# Pre- and post-synaptic roles for DCC in memory consolidation in the adult mouse hippocampus

**DOI:** 10.1101/697631

**Authors:** Stephen D. Glasgow, Edwin W. Wong, Greta Thompson-Steckel, Philippe Séguéla, Edward S. Ruthazer, Timothy E. Kennedy

**Affiliations:** Montréal Neurological Institute, Department of Neurology & Neurosurgery, 3801 Rue University, McGill University, Montréal, Québec, Canada H3A 2B4; NSERC CREATE Neuroengineering Training Program, McGill University

**Keywords:** CA1 pyramidal neurons, long-term potentiation, axon guidance, spatial memory

## Abstract

The receptor deleted in colorectal cancer (DCC) and its ligand netrin-1 are essential for axon guidance during development and are expressed by neurons in the mature brain. Netrin-1 recruits GluA1-containing α-amino-3-hydroxy-5-methyl-4-isoxazolepropionic acid receptors (AMPARs) and is critical for long-term potentiation (LTP) at CA3-CA1 hippocampal Schaffer collateral synapses, while conditional DCC deletion from glutamatergic neurons impairs hippocampal-dependent spatial memory and severely disrupts LTP induction. DCC co-fractionates with the detergent-resistant component of the postsynaptic density, yet is enriched in axonal growth cones that differentiate into presynaptic terminals during development. Specific presynaptic and postsynaptic contributions of DCC to the function of mature neural circuits have yet to be identified. Employing hippocampal subregion-specific conditional deletion of DCC, we show that DCC loss from CA1 hippocampal pyramidal neurons results in deficits in spatial memory, increased resting membrane potential, abnormal dendritic spine morphology, and weaker spontaneous excitatory postsynaptic activity. In contrast, deletion of DCC from CA3 neurons did not induce detectable changes in spine morphology or intrinsic electrophysiological properties of CA1 pyramidal neurons, but resulted in impaired performance on the novel object place recognition task as well as compromised excitatory synaptic transmission and long-term potentiation (LTP) at the Schaffer collateral synapse. Together, these findings reveal that DCC makes specific pre- and post-synaptic contributions to hippocampal synaptic plasticity underlying spatial memory.

## Introduction

Long-term potentiation (LTP) is an extensively studied form of activity-dependent synaptic plasticity (Bliss & Collingridge, 1993). Brief high-frequency stimulation (HFS) of Schaffer collateral synapses, the primary excitatory efferent connection between CA3 and CA1 pyramidal neurons in the hippocampus, results in a long-lasting change in the strength of synaptic transmission that is mediated primarily through the modification and membrane recruitment of excitatory postsynaptic receptors on dendritic spines (Diering & Huganir, 2018; Lamprecht & LeDoux, 2004). Accordingly, alteration of dendritic spine structure and synaptic composition can impact neuronal transmission and activity-dependent plasticity.

First described in the embryonic central nervous system, the secreted chemotropic guidance cue, netrin-1, regulates cytoskeletal reorganization, mediates cell adhesion, and directs cell and axon migration (Keino-Masu et al., 1996; Kennedy, Serafini, de la Torre, & Tessier-Lavigne, 1994; Lai Wing Sun, Correia, & Kennedy, 2011). In the mature nervous system, conditional deletion of netrin-1 from principal excitatory forebrain neurons impairs activity-dependent plasticity, and bath application of exogenous netrin-1 results in rapid synaptic recruitment of Ca^2+^-permeable AMPARs (Glasgow et al., 2018). Further, hippocampal-dependent spatial memory is impaired in mice conditionally lacking netrin-1 in principal excitatory forebrain neurons (Wong et al., 2019), suggesting that netrin-1 signaling has a critical role in synaptic plasticity underlying hippocampal-dependent spatial memory.

The canonical netrin-1 receptor, deleted in colorectal cancer (DCC), is essential for neural development and widely expressed in the mature nervous system (Keino-Masu et al., 1996; Volenec, Bhogal, Moorman, Leslie, & Flanigan, 1997). DCC null mice die within a few hours following birth, making it impossible to study DCC loss-of-function in adults using conventional knockouts (Fazeli et al., 1997). To investigate the specific functional role of DCC at presynaptic and postsynaptic sides of the Schaffer collateral synapse in the mature brain, we removed a floxed *dcc* allele using subregion-specific Cre recombinase driver mouse lines. Selective deletion of DCC from CA1 glutamatergic neurons increased thin-type dendritic spines with concomitant reduction in mushroom-type dendritic spines, decreased amplitude of spontaneous excitatory postsynaptic currents, and impaired spatial memory, but resulted in no change in HFS-induced LTP. In contrast, mice selectively lacking DCC expression by CA3 glutamatergic neurons showed impaired contextual spatial memory accompanied by reduced basal excitatory synaptic transmission and attenuated HFS-induced LTP; our findings provide evidence that these deficits are due to reduced DCC-mediated presynaptic vesicular mobilization in CA3 neurons. These findings reveal distinct pre- and postsynaptic functions for DCC at Schaffer collateral synapses contributing to synaptic plasticity underlying memory consolidation in the adult brain.

## Results

### Subregion-specific conditional deletion of DCC in the adult hippocampus

DCC is enriched at synapses and widely expressed by neurons throughout the adult brain, including the hippocampus (Horn et al., 2013; Volenec et al., 1997). To investigate whether DCC contributes to synaptic transmission in the adult brain through distinct pre- and post-synaptic mechanisms, we selectively deleted a floxed *dcc* allele from either CA3 or CA1 excitatory pyramidal neurons by Cre expression regulated by Grik4 (Grik4-Cre/DCC^*fl/fl*^) or R4ag11 (R4ag11-Cre/DCC^*fl/fl*^) promoters, respectively. Cre recombinase in Grik4-Cre and R4ag11-Cre mice is first detectable at P14 and P17, respectively, and recombination is maximal at 8 weeks of age (Dragatsis & Zeitlin, 2000; Nakazawa et al., 2002; Zakharenko et al., 2003).

Critically, Cre expression is initiated after developmental axon guidance is complete, and both Grik4-Cre/DCC^*fl/fl*^ and R4ag11-Cre/DCC^*fl/fl*^ mice are viable. To confirm hippocampal subregion selectivity of DCC deletion, we assessed levels of DCC protein using Western blot analyses. Homogenates of microdissected adult hippocampi revealed reduced levels of DCC protein in CA1 homogenate from R4ag11-Cre/DCC^*fl/fl*^ mice compared to Cre-negative DCC^*fl/fl*^ wildtype littermates (Figure 1A). Levels of DCC protein were comparable in isolated CA3 homogenates, indicating specific deletion of DCC from CA1 pyramidal neurons. Equal levels of DCC protein were detected in microdissected CA1 homogenates in Grik4-Cre/DCC^*fl/fl*^ mice and Cre-negative DCC^*fl/fl*^ littermates. In contrast, less DCC protein was detected in isolated CA3 homogenates from Grik4-Cre/DCC^*fl/fl*^ mice compared to wild-type littermates, consistent with specific deletion of DCC from CA3 pyramidal neurons using Grik4-Cre/DCC^*fl/fl*^ mice (Figure 1A). These findings confirm subregion-specific deletion of DCC in the mature hippocampus using targeted Cre recombination.

**Figure 1.**
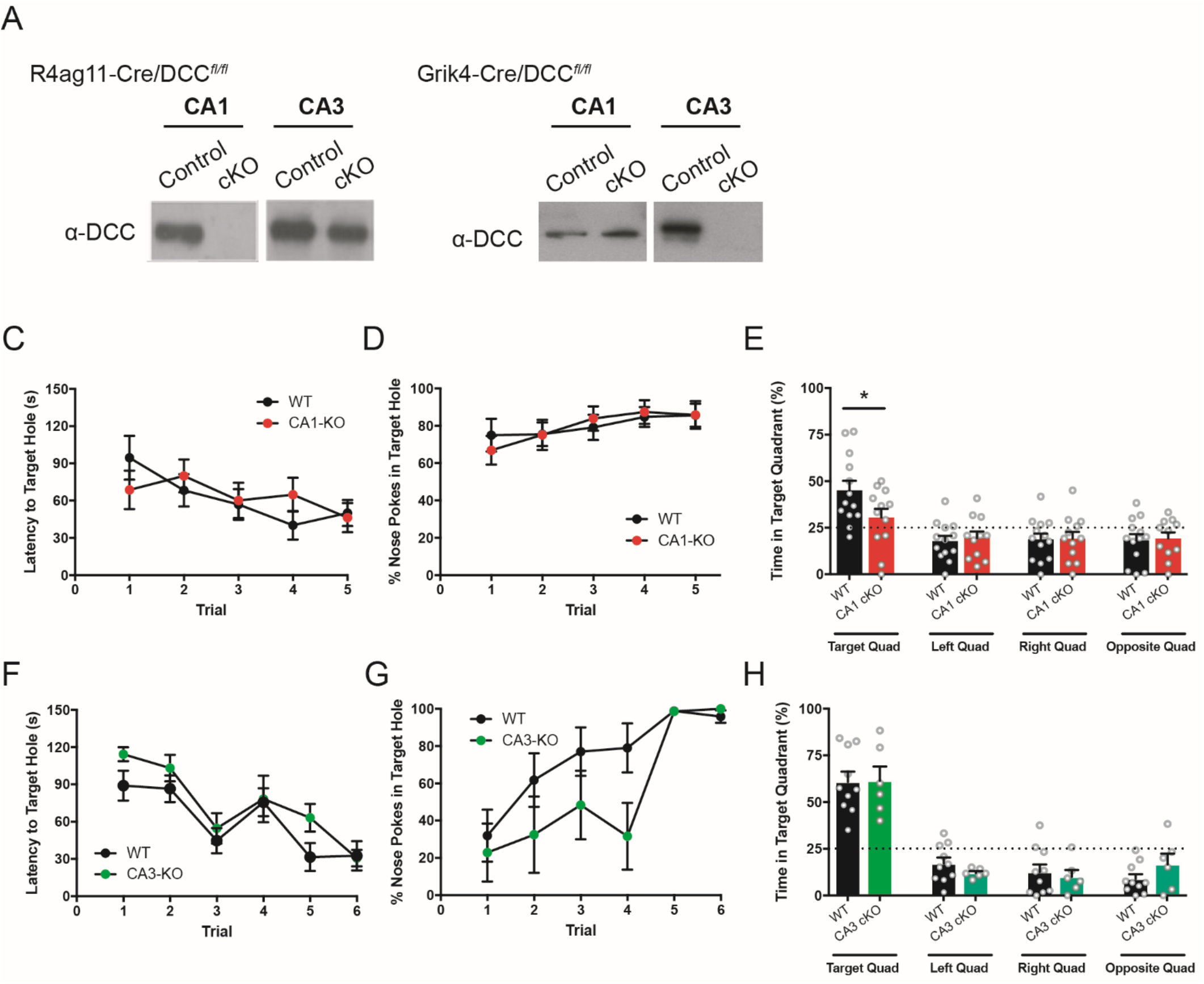
Conditional deletion of DCC in CA1 pyramidal hippocampal neurons but not CA3 pyramidal hippocampal neurons impairs spatial memory performance on Barnes maze. (**A**) Western blots of microdissected CA1 homogenate (left) and CA3 homogenate (right) from wild-type littermates and R4ag11 CamKIIα-Cre/DCC^*f/f*^ mice and Grik4-Cre/DCC^*f/f*^ mice. (**B-D**) R4ag11-Cre/DCC^*fl/fl*^ (CA1 cKO) show no differences in latency to (**B**; 0.36) nor number of nose pokes in target hole (**C**; *F*_5,70_=1.32, *p*=0.26) during training phase, but spent significantly less time in the target hole quadrant compared to control littermates (black) (**D**; R4ag11-Cre/DCC^*f/f*^: 30.5±4.7 s, Control: 45.0±5.2 s; *t*_23_=2.09, *p*=0.04). (**E-G**) Grik4-Cre/DCC^*fl/fl*^ (CA3 cKO) did not show any significant differences in escape latency (**E**) or nose pokes into target hole (**F**; Interaction: *F*_4,56_ = 0.96, *p* = 0.43; Main effect of genotype: *F*_1,56_ = 4.36, *p* = 0.055; Main effect of time: *F*_4,56_ = 7.67, *p*<0.001)) during training, and did not differ from control mice in time spent in target quadrant (**G**; *t*_14_=0.05, *p*=0.96). * denotes *p*<0.05.

### Selective deletion of DCC from CA1 glutamatergic neurons impairs spatial memory

In a previous study, DCC co-fractionated with detergent-resistant components of postsynaptic density, and both confocal and immuno-electron microscopy supported post-synaptic enrichment of DCC, however, these findings did not rule out the possibility that DCC might be also be present on the presynaptic side of the Schaffer collateral synapse (Horn et al., 2013). DCC expression by glutamatergic excitatory neurons is required for spatial memory formation (Horn et al., 2013), however DCC was deleted from both CA1 and CA3 hippocampal pyramidal neurons in these studies and respective functional contributions made by presynaptic and postsynaptic DCC were not addressed.

To assess the role of CA1 DCC expression in the consolidation of hippocampal-dependent spatial memory, R4ag11-Cre/DCC^*fl/fl*^ and wild-type age-matched littermate mice (Cre-negative DCC^*fl/fl*^ mice) were first tested on a modified Barnes maze paradigm. Briefly, mice were trained over 2 days on a circular open field table with equally spaced holes. One hole (“target”) was attached to an escape box, and relies on the innate preference of rodents to seek enclosed spaces compared to open fields (Attar et al., 2013; Barnes, 1979). R4ag11-Cre/DCC^*fl/fl*^ mice did not exhibit any difference in the latency to escape to (Figure 1C), or number of nose pokes (Figure 1D) into the target hole, compared to control mice, indicating intact sensory and motor functions, and consistent with normal cognitive spatial map formation (Brody & Holtzman, 2006). Twenty four hrs following the final training session, the mice were subjected to a probe trial in which the escape box was removed. R4ag11-Cre/DCC^*fl/fl*^ mice spent significantly less time in the target quadrant compared to control littermates, and explored each quadrant at chance levels (25%) (Figure 1E). These findings provide evidence that deletion of DCC from CA1 pyramidal neurons results in a significant impairment in spatial memory performance.

Previous work has suggested that DCC is localized to the presynaptic terminal of developing retinal ganglion cell (RGC) axons during target innervation (Manitt, Nikolakopoulou, Almario, Nguyen, & Cohen-Cory, 2009), and although substantial evidence supports post-synaptic DCC localization and function, presynaptic DCC at the Schaffer collateral synapse had not been ruled out. To determine a role for DCC expression in CA3 pyramidal neurons during spatial memory, we tested the performance of Grik4-Cre/DCC^*fl/fl*^ mice and their control wild-type littermates on the same modified Barnes maze. No significant differences were detected in escape latency (Figure 1F), and Grik4-Cre/DCC^*fl/fl*^ mice showed a non-significant decrease in the number of nose pokes into the target hole (Figure 1G) during the training phase (*p* = 0.055). Following a 24 hr delay, mice were assessed for time spent in the quadrant previously containing the escape box. In contrast to selective deletion of DCC from CA1 pyramidal neurons, Grik4-Cre/DCC^*fl/fl*^ mice spent significantly more time in the target quadrant compared to other non-target quadrants, and did not differ significantly compared to wild-type littermates (Figure 1H). These findings indicate that selective deletion of DCC from CA3 pyramidal neurons does not impair spatial memory performance in the Barnes maze.

Impaired Barnes maze performance suggests a deficit in spatial memory function in mice lacking DCC expressed by CA1 hippocampal pyramidal neurons, however previous reports suggest that Barnes maze protocols may be anxiogenic (Pitts, 2018). To assess whether mice conditionally-lacking DCC in CA1 or CA3 pyramidal neuron showed impaired hippocampal-dependent contextual spatial memory function using a non-anxiogenic task, we assessed R4ag11-Cre/DCC^*fl/fl*^ or Grik4-Cre/DCC^*fl/fl*^ mice using a novel object place recognition test (NOPR; Figure 2A) (Boyce, Glasgow, Williams, & Adamantidis, 2016; Wong et al., 2019). During the test phase, we observed no significant differences in total exploration time between R4ag11-Cre/DCC^*fl/fl*^ and control mice (Figure 2B). However, consistent with impaired spatial memory, R4ag11-Cre/DCC^*fl/fl*^ mice spent significantly less time investigating the displaced object (Figure 2C) and showed a significant reduction in investigation ratio compared to wild-type littermates (Figure 2D). Grik4-Cre/DCC^*fl/fl*^ mice showed no difference in total exploration time (Figure 2E), but spent less time with the displaced object (Figure 2F) and showed decreased investigation ratio compared to control mice (Figure 2G), indicating that DCC deletion from CA3 neurons impairs spatial memory performance on the NOPR task. These findings provide evidence that DCC expression by both CA1 and CA3 pyramidal neurons contribute to contextual spatial memory formation that requires object discrimination.

**Figure 2.**
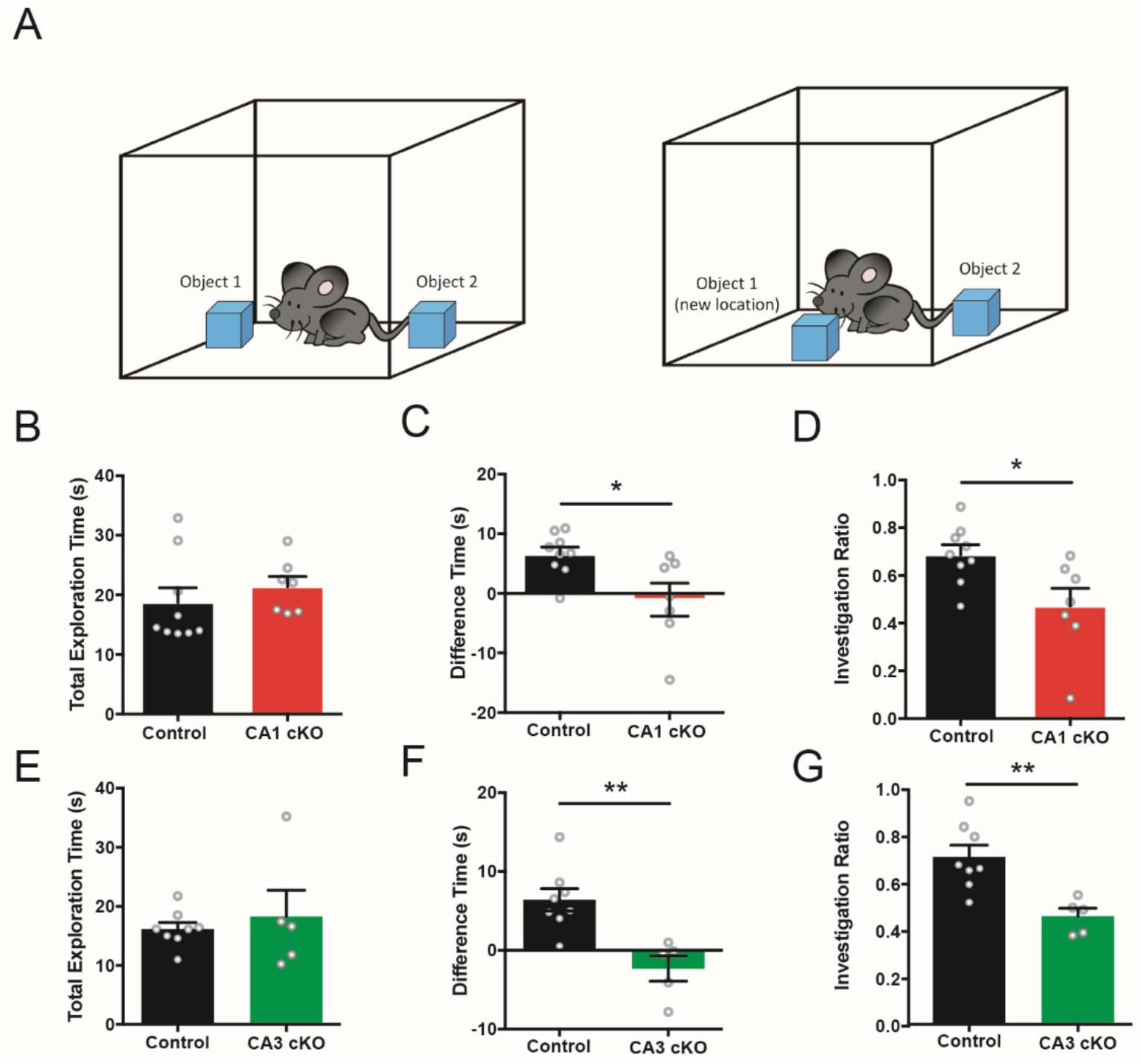
Conditional deletion of DCC from CA1 and CA3 pyramidal neurons impairs novel object place recognition. (**A**) Schematic representation of the novel object place recognition task. (**B-D**) R4ag11-Cre/DCC^*fl/fl*^ mice did not show significant differences in the total exploration time for either object (**B**; R4ag11-Cre/DCC^*fl/fl*^: *n* = 7, 21.4±1.7 s,, Control: *n* = 9, 18.7±2.5 s; *p* = 0.42), but spent significantly less time with the displaced object (**C**; R4ag11-Cre/DCC^*fl/fl*^: *n* = 7, −1.0±2.8 s,, Control: *n* = 9, 6.5±1.2 s; *t*_14_=2.74, *p* = 0.016) and showed reduced investigation ratio (**D**; R4ag11-Cre/DCC^*fl/fl*^: *n* = 7, 0.47±0.08, Control: *n* = 9, 0.69±0.04; *t*_14_=2.69, *p* = 0.017). (**E-G**) Conditional deletion of DCC from CA3 pyramidal neurons resulted in no differences in total exploration time (E; Grik4-Cre/DCC^*fl/fl*^: *n*=5, 18.3±4.5 s, Control: *n*=8, 16.2±1.1 s, p=0.59) compared to control mice (black), but significantly less time was spent with the displaced object (F; Grik4-Cre/DCC^*fl/fl*^: *n*=5, −2.3±1.6 s, Control: *n*=8, 6.4±1.4 s; *t*_11_=3.93, *p*=0.002) and the investigation ratio was showed significantly reduced (G; Grik4-Cre/DCC^*fl/fl*^: *n*=5, 0.46±0.03, Control: *n*=8, 0.71±0.05; *t*_11_=3.66, *p*=0.003). * denotes *p*<0.05.

Recognition of object location novelty evaluates the ability to recognize a familiar object in a new location, and co-opts intrinsic novelty-seeking behaviour (Antunes & Biala, 2012). However, NOPR requires visual discrimination and intact object recognition, which are mediated by non-hippocampal brain regions (Mumby & Pinel, 1994). To examine whether non-hippocampal object recognition performance was impaired in mice conditionally-lacking DCC from either CA1 or CA3 pyramidal neurons, we assessed R4ag11-Cre/DCC^*fl/fl*^ or Grik4-Cre/DCC^*fl/fl*^ mice using a novel object recognition task (Figure 3A). Consistent with normal visual function, we observed no significant differences in the total exploration time, proportion of time spent exploring novel objects, or investigation ratio in R4ag11-Cre/DCC^*fl/fl*^ (Figure 3B-D) or Grik4-Cre/DCC^*fl/fl*^ (Figure 3E-G) mice compared to control littermates. These findings indicate that mice lacking neuronal DCC expression in hippocampal subregions are not impaired in their ability to discriminate between novel visual cues.

**Figure 3.**
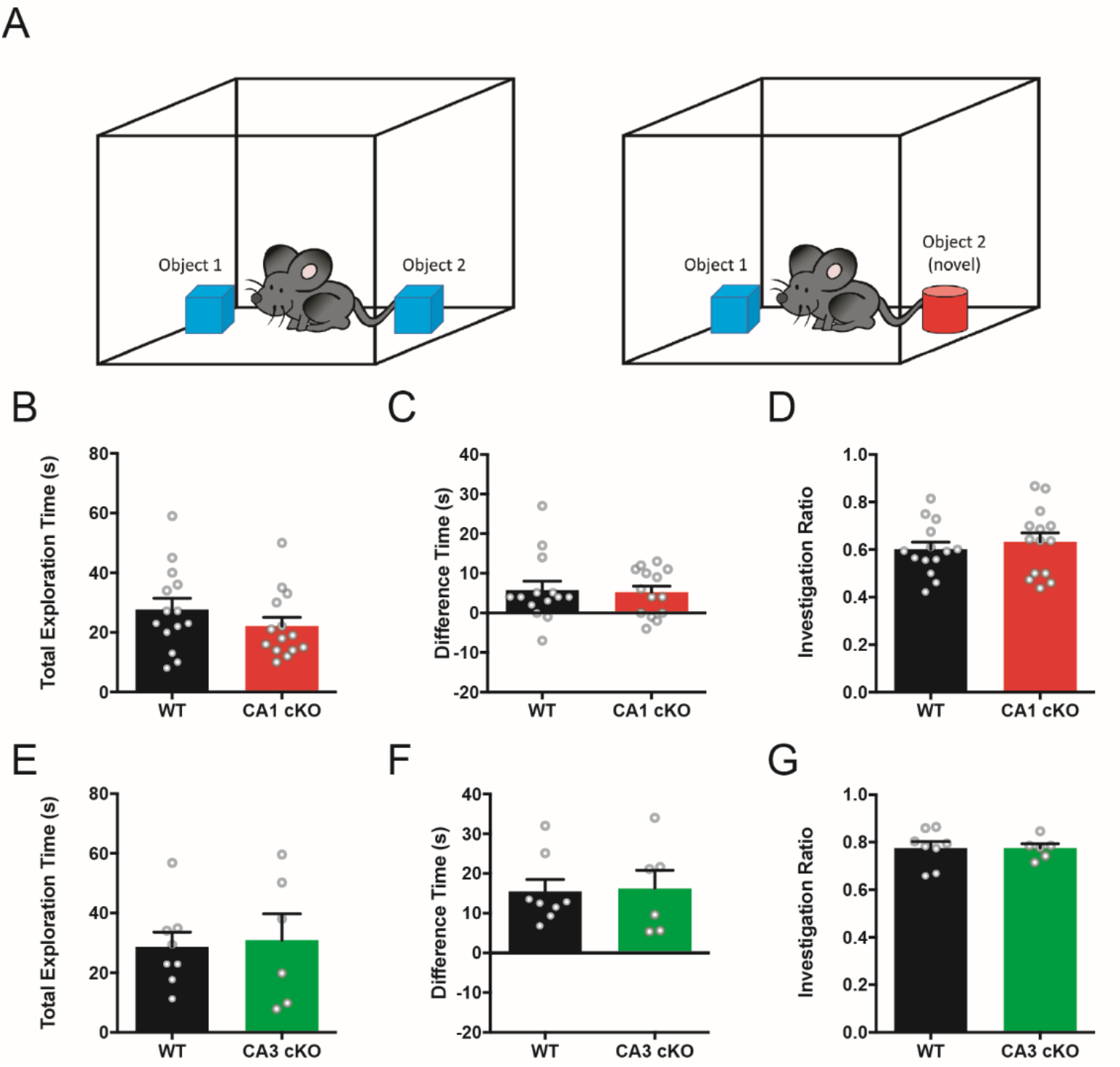
Conditional deletion of DCC from CA1 or CA3 pyramidal neurons does not impair performance on novel object recognition task. (**A**) Schematic representation of the novel object recognition task. (**B-D**) R4ag11-Cre/DCC^*fl/fl*^ (red) show no significant differences in total exploration time (**B**; R4ag11-Cre/DCC^*fl/fl*^: *n*=14, 22.1±3.0 s, Control: *n*=14, 27.6±3.7 s; *p*=0.26), time spent with novel object (**C**; R4ag11-Cre/DCC^*fl/fl*^: *n*=14, 5.2±1.6 s, Control: *n*=14, 5.8±2.3 s; *p*=0.83), or investigation ratio (**D**; R4ag11-Cre/DCC^*fl/fl*^: *n*=14, 0.63±0.4, Control: *n*=14, 0.60±0.03 s; *p*=0.52) compared to control littermates (black). (**E-G**) Grik4-Cre/DCC^*fl/fl*^ did not spend more total time exploring objects (**E**; Grik4-Cre/DCC^*fl/fl*^: *n*=6, 30.88 ± 8.851 s, Control: n=8, 28.7±4.9 s, *p*=0.82), time spent with novel object (**F**; Grik4-Cre/DCC^*fl/fl*^: *n*=6, 16.2 ± 4.6 s, Control: *n*=8, 15.5±3.0 s, *p*=0.89), or investigation ratio (**G**; Grik4-Cre/DCC^*fl/fl*^: *n*=6, 0.78 ± 0.02, Control: *n*=8, 0.78±0.02 s, *p*=0.99).

### Deletion of DCC does not impair hippocampal neuronal excitability

Changes in spatial memory function have been associated with alteration in overall neuronal excitability (Oh, Oliveira, & Disterhoft, 2010). To assess whether conditional deletion of DCC alters intrinsic cellular excitability of CA1 pyramidal neurons, we recorded voltage responses to intracellular current injections using whole-cell patch clamp recordings from CA1 pyramidal neurons in acute hippocampal brain slices derived from 8-10 month old R4ag11-Cre/DCC^*fl/fl*^ or Grik4-Cre/DCC^*fl/fl*^ mice (Figure 4A, D). Previous work provided evidence that DCC activates phospholipase C-γ (PLCγ) (Horn et al., 2013; Xie et al., 2006), which in turn can modulate the open probability of various K^+^ channels through reduction of membrane-bound phosphatidylinositol 4,5-bisphosphate (PIP_2_) and result in alteration of neuronal input resistance (Kobrinsky, Mirshahi, Zhang, Jin, & Logothetis, 2000). Interestingly, we observed no change in steady-state input resistance (Figure 4B), however we detected a slight hyperpolarization in the resting membrane potential of CA1 pyramidal neurons from R4ag11-Cre/DCC^*fl/fl*^ mice compared to control littermates (Figure 4C). In contrast, no significant alteration of intrinsic cellular properties was detected in CA1 pyramidal neurons from Grik4-Cre/DCC^*fl/fl*^ mice (Figure 4D-F).

**Figure 4.**
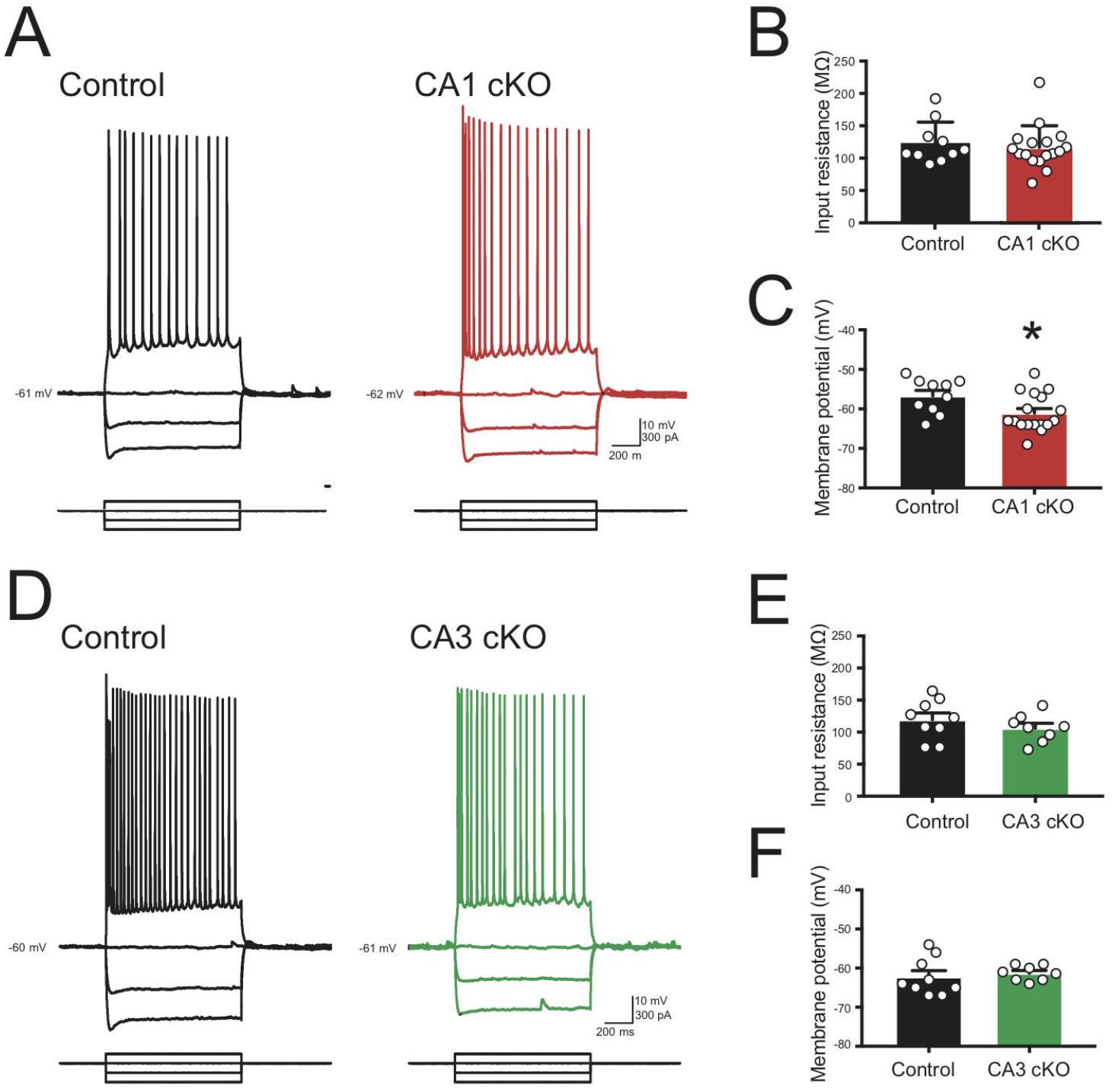
Intrinsic excitability of CA1 pyramidal neurons in R4ag11-Cre/DCC^*fl/fl*^ and Grik4-Cre/DCC^*fl/fl*^ mice. **(A)** Membrane potential responses to hyperpolarizing and depolarizing current pulses in CA1 pyramidal neurons derived from age-matched control (black, left) and R4ag11-Cre/DCC^*fl/fl*^ (CA1 cKO; red, right). (**B-C**) Group data showing resting membrane potential (**B**; R4ag11-Cre/DCC^*fl/fl*^: −56.6±1.5 mV, Control: −61.1±1.1 mV, *t*_25_=2.39, *p*=0.024) and input resistance (**C**; R4ag11-Cre/DCC^*fl/fl*^: 116±8 MΩ, Control: 123±10 MΩ, *p*=0.602) for CA1 pyramidal neurons from R4ag11-Cre/DCC^*fl/fl*^ and control mice. (**D**) Membrane potential traces in response to hyperpolarizing and depolarizing current pulses in CA1 pyramidal neurons from control (black, left) and Grik4-Cre/DCC^*fl/fl*^ (CA3 cKO; green, right). (**E-F**) Group data show no significant differences in input resistance (**E**; Grik4-Cre/DCC^*fl/fl*^: 119±10 MΩ, Control: 106±7 MΩ, *p*=0.325) or resting membrane potential (**F**; Grik4-Cre/DCC^*fl/fl*^: −61.3±0.7 mV, Control: −62.2±1.6 mV, *p*=0.62) between Grik4-Cre/DCC^*fl/fl*^ mice and control littermates. * denote *p*<0.05.

### Effect of pre- and post-synaptic deletion of DCC on the Schaffer collateral synapse

We have previously demonstrated that conditional deletion of DCC from principal excitatory neurons in the forebrain and hippocampus can strongly attenuate LTP at Schaffer collateral synapses in the adult hippocampus (Horn et al., 2013). While substantial evidence indicates that DCC is present post-synaptically, pre-synaptic DCC had not been ruled out and it was not clear if pre- or post-synaptic DCC function is required for synaptic plasticity in the adult brain. To assess the effect of conditional deletion of DCC from either pre- or postsynaptic neurons in the hippocampus, we first recorded excitatory postsynaptic responses (EPSCs) from CA1 pyramidal neurons in response to stimulation of CA3 Schaffer collateral axons. Basal evoked EPSCs were measured in response to pulses presented with 25 μA increments in stimulation intensity up to 200 μA (Figure 5A). No appreciable changes in amplitude of synaptic responses (Figure 5B) or paired-pulse ratios (Figure 5C; 10-200 ms ISI) were detected between R4ag11-Cre/DCC^*fl/fl*^ and wild-type controls. In contrast, the amplitude of evoked EPSCs in CA1 pyramidal neurons in response to Schaffer collateral stimulation was significantly lower at high stimulation intensities (150-200 μA) in Grik4-Cre/DCC^*fl/fl*^ mice compared to control littermates (Figure 5D-E). These findings indicate that deletion of DCC from presynaptic CA3 neurons results in significantly decreased excitatory synaptic drive onto CA1 pyramidal neurons in the adult hippocampus, and suggest that DCC may influence the dynamics of presynaptic release.

**Figure 5.**
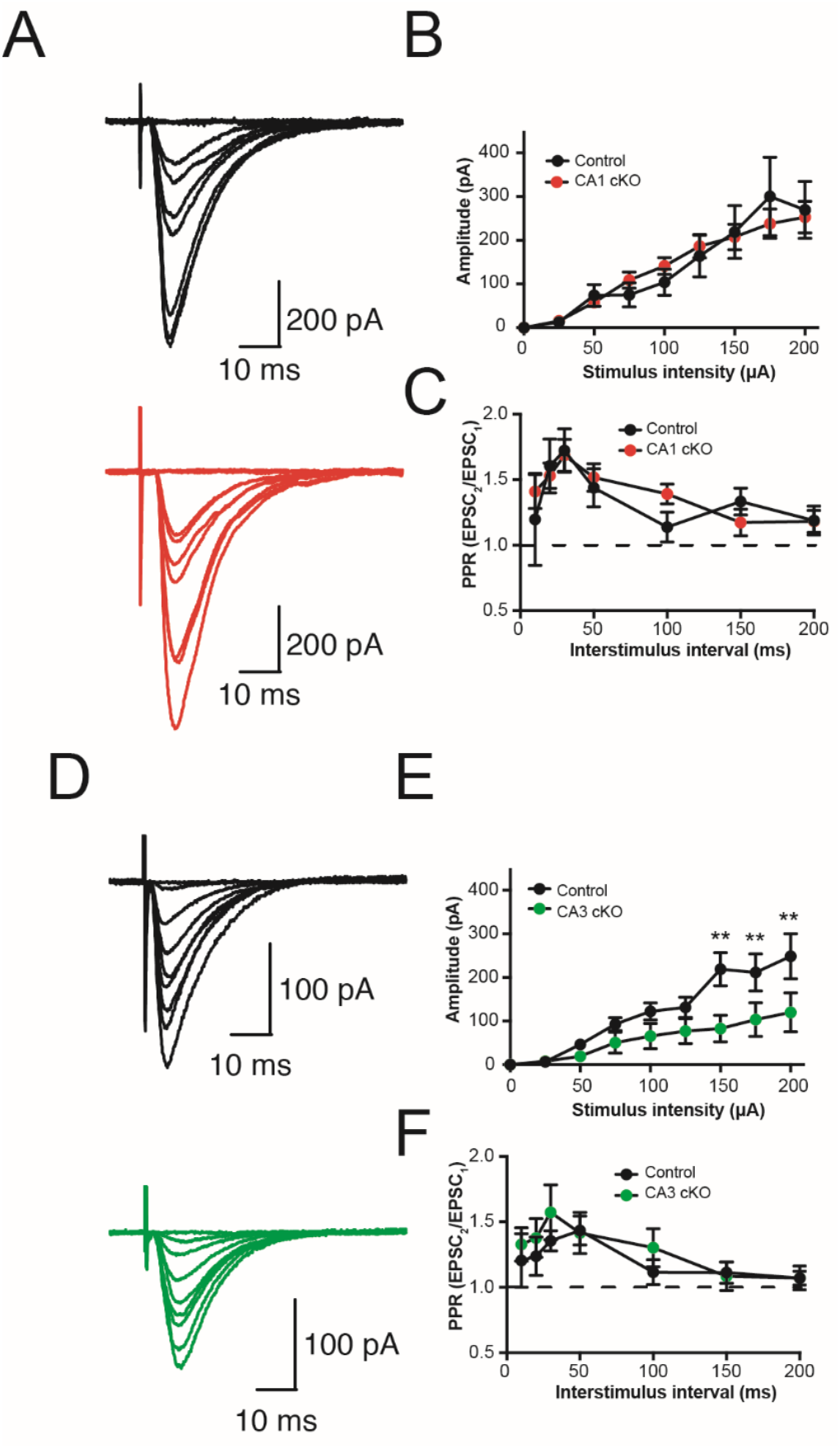
Selective deletion of DCC from CA3 pyramidal neurons impairs evoked synaptic transmission at Schaffer collateral synapses. **(A)** Representative traces of synaptic responses in CA1 pyramidal neurons in response to Schaffer collateral stimulation from control wild-type littermates (top, black) and R4ag11-Cre/DCC^*fl/fl*^ (CA1 cKO; bottom, red). (**B-C**) Group data show evoked EPSC amplitude in CA1 pyramidal neurons in response to incremental stimulation intensity of Schaffer collaterals (**B**; *F*_8,192_=0.18, *p*=0.993) and paired-pulse ratio across a range of interstimulus intervals (**C**; ISI; Interaction: *F*_6,78_=0.76, *p*=0.602) in R4ag11-Cre/DCC^*fl/fl*^ (CA1 cKO, red) and control littermates (black). (**D-F**) Representative traces (**D**) and group data (**E**; Interaction: *F*_8,112_=3.13, *p*=0.003) from CA1 pyramidal neurons in control wild-type littermates (top, black) and Grik4-Cre/DCC^*fl/fl*^ (CA3 cKO; bottom, green) in response to incremental stimulation intensity of Schaffer collateral. (**F**) Group data also show paired-pulse ratio across a range of ISI (Interaction: *F*_6,84_=0.53, *p*=0.780). ** denotes *p*<0.01.

DCC has been reported to associate with TRIM9, which can modulate soluble N-ethylmaleimide-sensitive factor-attachment protein (SNAP) receptor (SNARE)-mediated exocytosis mechanisms to regulate vesicular trafficking and neurotransmitter release (Plooster et al., 2017; Winkle et al., 2014). To determine whether conditional deletion of DCC was associated with changes in presynaptic release dynamics, we assessed the paired pulse ratio across a range of interstimulus intervals (10-200 ms ISI) in Grik4-Cre/DCC^*fl/fl*^ mice and control littermates. No change in PPR was detected across all ISIs in Grik4-Cre/DCC^*fl/fl*^, consistent with a lack of effect of either pre- or postsynaptic DCC deletion on the probability of presynaptic vesicular release (Figure 5F).

Deletion of DCC from CA3 and CA1 pyramidal neurons (Horn et al., 2013) or selective conditional deletion from either CA1 pyramidal neurons (R4ag11-Cre/DCC^*fl/fl*^) or CA3 pyramidal neurons (Grik4-Cre/DCC^*fl/fl*^) impairs contextual spatial memory performance (Figure 2). Spatial memory performance requires *de novo* formation of location-specific neuronal firing fields (place fields), and place cell firing patterns can induce long-lasting NMDAR-dependent changes in the strength of synaptic connections, suggesting that deletion of DCC may disrupt synaptic potentiation in response to high-frequency stimulation (Isaac, Buchanan, Muller, & Mellor, 2009). To determine if postsynaptic deletion of DCC impairs LTP, we recorded evoked EPSCs in CA1 pyramidal neurons from R4ag11-Cre/DCC^*fl/fl*^ mice in response to Schaffer collateral stimulation in the presence of the GABA_A_ antagonist, picrotoxin (100 μM) (Figure 6A). Brief high-frequency stimulation (100 Hz for 1 s) resulted in significant short-term facilitation of EPSCs in both R4ag11-Cre/DCC^*fl/fl*^ mice and control littermates. Surprisingly, the amplitude of EPSCs remained significantly potentiated after 25 min in slices from both R4ag11-CreiDCC^*fl/fl*^ (144±10% of baseline, *p*<0.001) and controls (140±12% of baseline, *p*=0.02) compared to baseline values (Figure 6B-C). Furthermore, no change in paired-pulse ratio occurred (Figure 6D). These findings demonstrate that conditional postsynaptic deletion of DCC from CA1 pyramidal neurons does not impact NMDAR-dependent LTP at Schaffer collateral synapses.

Although primarily mediated through changes in the postsynaptic neuron, LTP may also involve alteration of pre-synaptic function, and homeostatic changes in release dynamics may contribute to Hebbian synaptic plasticity (Emptage, Reid, Fine, & Bliss, 2003; Soares, Lee, & Beique, 2017). To determine whether deletion of DCC from CA3 pyramidal neurons might impair HFS-induced LTP at Schaffer collateral synapses, we recorded evoked EPSCs in CA1 pyramidal neurons derived from Grik4-Cre/DCC^*fl/fl*^ mice in response to Schaffer collateral stimulation (Figure 6E). A brief high-frequency tetanus (100 Hz for 1 s) resulted in an immediate potentiation of synaptic responses in CA1 pyramidal neurons. However, by 25 mins after HFS, synaptic responses in slices from Grik4-Cre/DCC^*fl/fl*^ returned to baseline values (113±6% of baseline values, *p*=0.913) compared to controls (145±9% of baseline, *p*=0.002) (Figure 6F-G). We observed no changes in PPR following HFS in either Grik4-Cre/DCC^*fl/fl*^ mice or their control littermates (Figure 6H). Together, these results indicate that deletion of presynaptic, but not postsynaptic, DCC impairs HFS-induced LTP in the Schaffer collateral pathway of the adult hippocampus.

**Figure 6.**
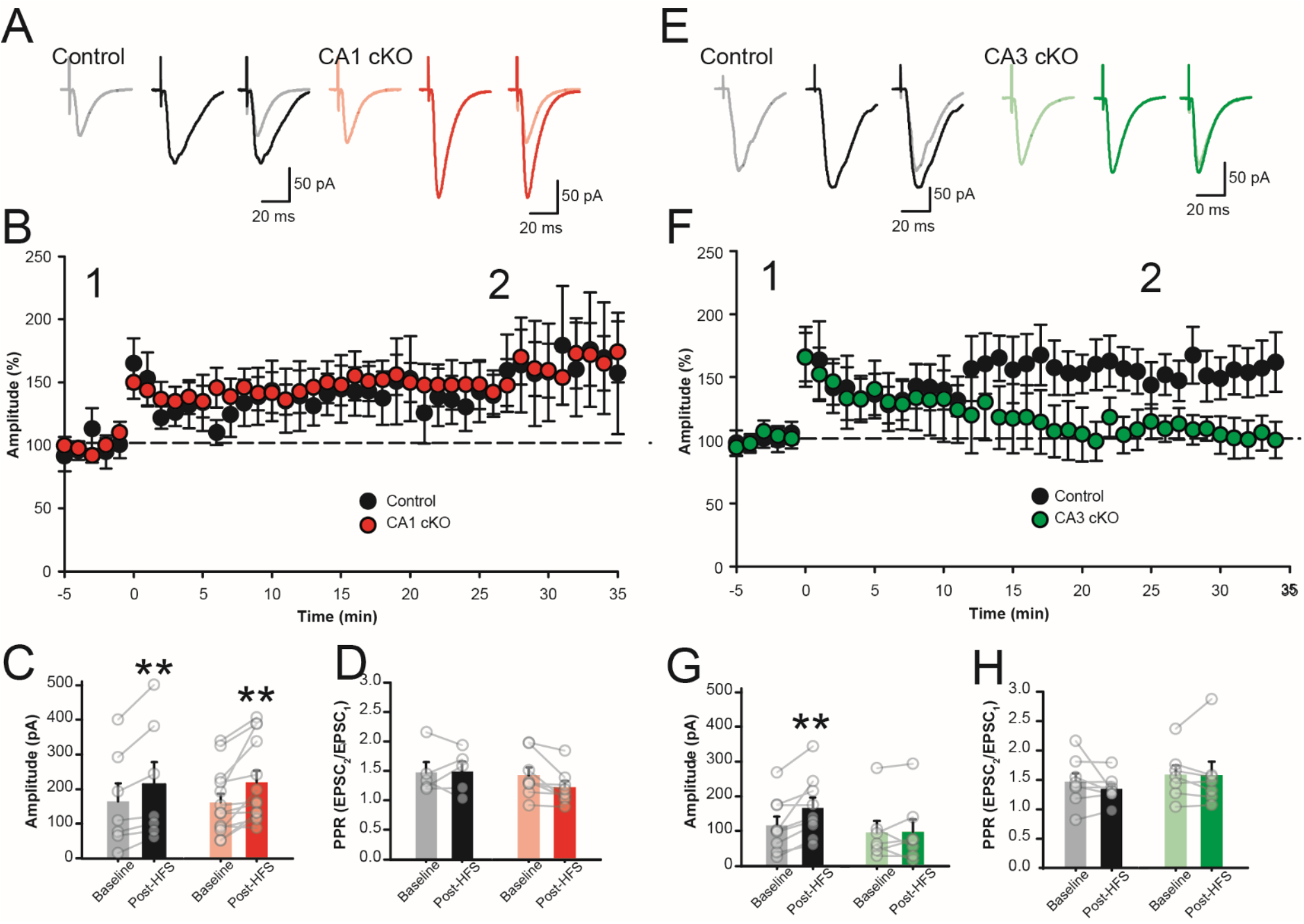
Deletion of DCC from CA3 pyramidal neurons, but not CA1 pyramidal neurons, impairs HFS-induced LTP in the adult hippocampus. **(A-B)** Representative evoked AMPAR-mediated EPSCs (A) and group data (B) recorded at −70 mV in response to Schaffer collateral stimulation from control littermates (left, black) and R4ag11-Cre/DCC^*fl/fl*^ (CA1 cKO; right, red) prior to and following (20 min post-HFS) brief high-frequency stimulation (1 s at 100 Hz). The amplitude of EPSCs remained significantly potentiated after 25 min in slices from both R4ag11-Cre/DCC^*fl/fl*^ (144±10% of baseline, *p*<0.001) and controls (140±12% of baseline, *p*=0.02) compared to baseline values (Main effect of HFS: *F*_1,18_=20.78, p<0.001; Interaction genotype X HFS: *F*_1,18_=0.20, *p*=0.659). (C) Paired-pulse ratio (50 ms ISI) was not significantly changed between R4ag11-Cre/DCC^*fl/fl*^ (CA1 cKO) mice and control littermates (*p*=0.602). (D-F) Representative evoked AMPAR-mediated EPSCs (D) and group data (E) recorded at −70 mV in response to Schaffer collateral stimulation from control littermates (left, black) and Grik4-Cre/DCC^*fl/fl*^ (CA3 cKO; right, green) prior to and following (20 min post-HFS) brief high-frequency stimulation (1 s at 100 Hz). AMPAR-mediated EPSCs were not significantly different from baseline values (113±6% of baseline values, *p*=0.913) compared to controls (145±9% of baseline, *p*=0.002) (Interaction for genotype X HFS: *F*_1,14_=6.30, *p*=0.025), with no appreciable change in the paired-pulse ratio (F; Interaction for genotype X HFS: *p*=0.780). ** denotes *p*<0.01.

### Postsynaptic deletion of DCC regulates basal synapse strength

While no differences in evoked synaptic strength were observed in R4ag11-Cre/DCC^*fl/fl*^ mice, we postulated that individual excitatory synapses may show reductions in sensitivity to presynaptic transmitter release. To assess basal synaptic strength in R4ag11-Cre/DCC^*fl/fl*^ mice, we recorded AMPAR-mediated spontaneous EPSCs (sEPSCs) in the presence of picrotoxin (100 μM) (Figure 7A). We found that the amplitude of sEPSC events were significantly decreased compared to events from age-matched control mice (Figure 7B). In contrast, we detected no significant differences in sEPSC frequency (Figure 7C). These findings indicate that deletion of DCC from CA1 pyramidal neurons impairs the relative strength of individual synapses, without affecting the overall number of synaptic contacts.

**Figure 7.**
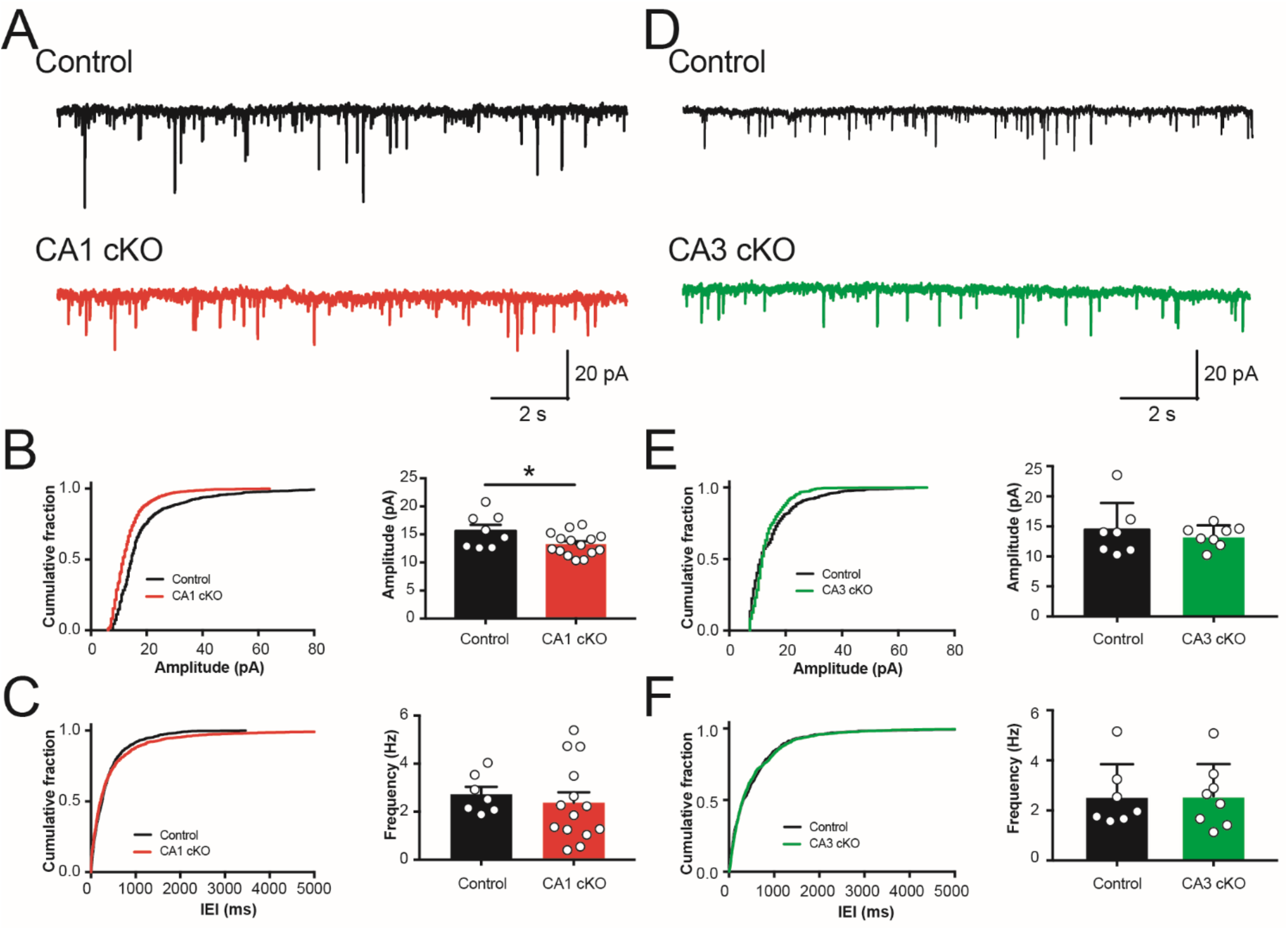
Conditional deletion of DCC from CA1 pyramidal neurons decreases sEPSC amplitude. (**A**) Representative spontaneous excitatory postsynaptic current (sEPSC) traces in CA1 pyramidal neurons from control (black, top) and R4ag11-Cre/DCCfl/fl (CA1 cKO; red, bottom). (**B-C**) Cumulative distribution plots (left) and group data (right) show a significant decrease in the average amplitude of sEPSCs in R4ag11-Cre/DCC^*fl/fl*^ mice (**B**; R4ag11-Cre/DCC^*fl/fl*^: *n*=15, 13.2±0.5 pA, Control: *n*=8, 15.6±1.0 pA; *t*_21_=2.26, *p* = 0.034) but no change in average frequency (**C**; R4ag11-Cre/DCC^*fl/fl*^: *n*=15, 2.3±0.4 Hz, Control: *n*=8, 2.7±0.3 Hz; *p*=0.59). (**D**) Representative sEPSC traces from CA1 pyramidal neurons from control (black, top) and Grik4-Cre/DCC^*fl/fl*^ mice (CA3 cKO; green, bottom). (**E-F**) Cumulative distribution plots (left) and group data (right) show no changes in average sEPSC amplitude (**E**; Grik4-Cre/DCC^*fl/fl*^: *n*=8, 13.3±0.6 pA, Control: *n*=7, 14.3±1.7 pA; *p*=0.60) or average frequency (**F**; Grik4-Cre/DCC^*fl/fl*^: *n*=8, 2.6±0.5 Hz, Control: *n*=7, 2.6±0.5 Hz; *p*=0.982) between Grik4-Cre/DCC^*fl/fl*^ and control littermates. K-S test for cumulative distribution plots. * denotes *p*<0.05.

Presynaptic deletion of DCC can prevent expression of HFS-induced LTP (Figure 6), however it remained unclear whether this resulted in alteration of network-level synaptic properties. To examine whether deletion of DCC in CA3 pyramidal neurons affected basal spontaneous synaptic transmission, we recorded sEPSCs in CA1 pyramidal neurons from Grik4-Cre/DCC^*fl/fl*^ mice in the presence of picrotoxin (100 μM) (Figure 7D). We detected no difference in average sEPSC amplitude or frequency between genotypes (Figure 7E-F), indicating that deletion of DCC from CA3 does not impair basal network-level synaptic inputs onto CA1 pyramidal neurons.

### Deletion of postsynaptic DCC alters dendritic spine morphology

Synaptic modification during learning has been proposed to play a critical role in the consolidation of new memories (Bliss & Collingridge, 1993). We observed that animals lacking DCC in CA1 pyramidal neurons exhibited significant deficits in spatial memory, but did not show any alteration in the propensity for CA1 LTP induction (Figures 1 and 6). However, we also identified smaller sEPSC amplitudes in R4ag11-Cre/DCC^*fl/fl*^ mice, suggesting a subtle change in excitatory synaptic transmission in the absence of DCC, which may underlie spatial memory impairments. Indeed, spatial memory function has been associated with changes in dendritic spine morphology in CA1 pyramidal neurons (Nimchinsky, Sabatini, & Svoboda, 2002; Yasumatsu, Matsuzaki, Miyazaki, Noguchi, & Kasai, 2008; Yuste & Bonhoeffer, 2001), and conditional deletion of DCC from both CA1 and CA3 pyramidal neurons in T29-1 *CamKIIα-Cre*/DCC^*fl/fl*^ mice results in an increase in stubby-type dendritic spines along the apical dendrites of CA1 pyramidal neurons (Horn et al., 2013). We therefore quantified dendritic spine density and morphology in CA1 pyramidal neurons from R4ag11-Cre/DCC^*fl/fl*^ and control littermates using the Golgi-Cox staining method (Figure 8A). Dendritic spine density was unchanged between R4ag11-Cre/DCC^*fl/fl*^ and control littermates (Figure 8B). In contrast, the average length and width of dendritic spines were significantly reduced along the apical dendrites of CA1 pyramidal neurons from R4ag11-Cre/DCC^*fl/fl*^ mice compared to control littermates (Figure 8B). These results indicate that postsynaptic deletion of DCC from CA1 pyramidal neurons does not affect dendritic spine density, but reduces average spine length and width along the apical dendrites of CA1 pyramidal neurons.

**Figure 8.**
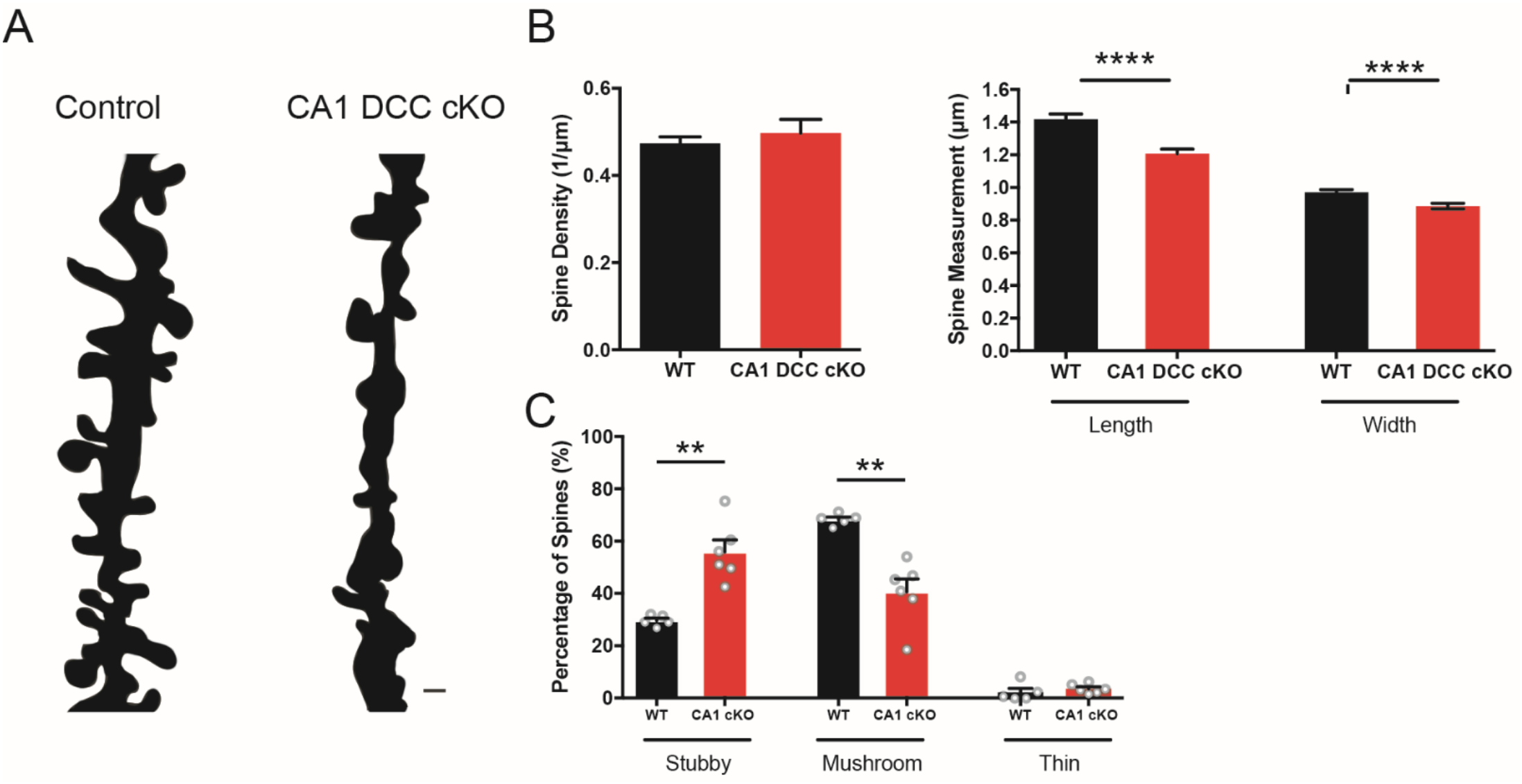
Alterations in dendritic spine morphology in CA1 pyramidal neurons from R4ag11-Cre/DCC^*fl/fl*^ mice. **(A)** Representative camera lucida reconstructions of CA1 pyramidal neuron apical dendrites from control (left) and R4ag11-Cre/DCC^*fl/fl*^ mice. Scale bar = 1 μm. (**B**) Group data show no significant difference in average spine density between control and R4ag11-Cre/DCC^*fl/fl*^ mice (left), but significant reductions in average spine length and spine head width (right). (**C**) Dendritic spine type classification reveals a significant increase in number of stubby-type spines (R4ag11-Cre/DCC^*fl/fl*^: 55.8±4.6%, Control: 29.6±0.9%; *t*_5.4_=5.59, *p*=0.002) and significant decrease in the number of mushroom-type dendritic spines (R4ag11-Cre/DCC^*fl/fl*^: 40.6±4.9%, Control: 68.2±1.0%; *t*_5.47_=5.40, *p*=0.002), with no detectable differences in thin-type spines (*p*=0.41). **: *p*<0.01. Data are presented as mean ± SEM.

Dendritic spines can be grouped into three major categories: thin, mushroom, and stubby (Rochefort & Konnerth, 2012). Thin-type spines have longer lengths than head diameters, and mushroom-type spines have head sizes that greatly exceed neck length (Bourne & Harris, 2007). Stubby-type spines are transient, characterized by short neck lengths and wide diameters, and are considered immature, showing robust increases in overall volume following plasticity-inducing protocols (Matsuzaki, Honkura, Ellis-Davies, & Kasai, 2004). To determine whether DCC deletion in CA1 pyramidal neurons impacted dendritic spine morphology, we examined the density of different spine types (mushroom, thin, stubby) in wild-type littermates and CA1 DCC cKO mice. We detected a significant increase in the number of stubby spines and reduction of mushroom spines in CA1 DCC cKO mice, with no significant change in thin spines (Figure 8C). Together, these findings identify a DCC-mediated contribution to the maturation, maintenance, and stabilization of mature spine structures in CA1 pyramidal neurons.

## Discussion

Recent evidence has demonstrated an emerging role for axon guidance proteins in the adult brain, however little is known regarding the locus of action for these molecules (Lai Wing Sun et al., 2011; Shen & Cowan, 2010; Tillo, Ruhrberg, & Mackenzie, 2012). Here, we show that pre- or post-synaptic deletion of the netrin-1 receptor, DCC, from excitatory glutamatergic neurons in the mouse hippocampus have complementary roles in spatial memory, synaptic transmission, and synaptic plasticity. Consistent with our previous work, we demonstrated that conditional deletion of DCC from CA1 pyramidal neurons resulted in significant impairment of spatial memory tasks, reduced sEPSC amplitude, and altered spine morphology compared to control wild-type littermates. Conditional deletion of DCC from CA3 pyramidal neurons selectively impaired contextual spatial memory performance, and resulted in a striking attenuation of HFS-induced LTP. These findings show that distinct distributions of DCC at the Schaffer collateral synapse are necessary for spatial information processing, as well as functional and structural synaptic plasticity in the adult hippocampus.

### Postsynaptic deletion of DCC may regulate late-phase synaptic changes following memory formation

Deletion of DCC expressed by CA1 pyramidal neurons did not affect overall spine density, yet increased the proportion of stubby-type spines and decreased mushroom-type spines. Stubby-type spines (~ 1 μM in diameter) predominate during early postnatal life, and may represent an early stage of synaptogenesis (Harris, 1999). During the second postnatal week in mice, the majority of synapses formed are stubby-type protrusions from dendritic shafts, which in turn decrease in number by adulthood, when ~80% of dendritic spines have a mushroom-type structure (Harris, Jensen, & Tsao, 1992). During early development, DCC and netrin-1 promote glutamatergic synaptogenesis, directing the accumulation of presynaptic and postsynaptic proteins such as synaptophysin and PSD-95, respectively (Goldman et al., 2013). Cre expression in R4ag11-Cre mice starts at ~P17, following completion of axon guidance but during the process of spine maturation (Bourne & Harris, 2007; Dragatsis & Zeitlin, 2000). Postsynaptic loss of DCC in CA1 pyramidal neurons may impair the maturation of stubby spines into more stable mushroom spines, and thereby limit the accumulation of PSD-95 and other membrane-associated guanylate kinases-related proteins such as SAP97, which is necessary for netrin-1 mediated potentiation of hippocampal synapses (Glasgow et al., 2018; Goldman et al., 2013). As a result, the lack of DCC in CA1 pyramidal neurons may arrest spine maturation and attenuate ongoing basal synaptic transmission in adulthood, reducing the average strength of EPSCs in adult CA1 pyramidal neurons.

Dendritic spine shape and size can regulate synaptic transmission through modulation of electrical resistance (Araya, Vogels, & Yuste, 2014). Stubby-type spines are considered relatively transient, with only ~17% of all stubby spines in the hippocampus persisting following a 24 h period, and contain fewer postsynaptic proteins compared to more mature spine types (De Roo, Klauser, Mendez, Poglia, & Muller, 2008). Consistent with a role in learning and memory, spatial training decreased the number of stubby spines and increased the number of mushroom-type spines along the apical dendrites of CA1 pyramidal neurons, suggesting that stubby spines mature into more stable mushroom-type spines following consolidation (Mahmmoud et al., 2015). Selective deletion of DCC from CA1 pyramidal neurons may therefore allow for normal spatial behaviour but impair consolidation due to impaired maturation and stabilization of immature synapses. A preponderance of relatively immature PSDs in transient dendritic spines may contribute to altered basal spontaneous synaptic amplitude in animals lacking DCC in CA1 pyramidal neurons (R4ag11-Cre/DCC^*fl/fl*^) (Figure 7). However, the persistence of LTP in R4ag11-Cre/DCC^*fl/fl*^ mice (Figure 6) suggests that an increase in the proportion of stubby spines is sufficient for the early phases of activity-dependent synaptic potentiation.

Consistent with impaired long-term memory consolidation, we observed behavioural deficits in spatial tasks that required >24 hr delay periods between training and test probes. Although not assessed here, it is possible that the HFS-induced LTP produced in R4ag11-Cre/DCC^*fl/fl*^ mice may not exhibit the subsequent protein synthesis-dependent LTP late phase (Steward & Schuman, 2001). Recent work using *Aplysia* neurons demonstrated that local mRNA translation at synaptic sites can be initiated by netrin-1, suggesting that DCC may regulate protein synthesis following early phases of plasticity (Kim & Martin, 2015). We have previously demonstrated that bath application of netrin-1 rapidly recruits synaptic GluA1-containing AMPARs through a DCC-dependent mechanism (Glasgow et al., 2018), but the contribution of netrin-1 to late phase LTP in the mammalian brain is not clear. Consistent with roles in multiple phases of LTP, initial increases in Ca^2+^-permeable AMPARs facilitate rapid synaptic strengthening, which can be followed by GluA1-containing endocytosis and protein synthesis-dependent synaptic stabilization through insertion of GluA2-containing Ca^2+^-impermeable receptors (Diering & Huganir, 2018; Migues et al., 2016; Park et al., 2018; Zhou et al., 2018).

### The role of presynaptic DCC in synaptic transmission and plasticity

DCC plays an important role in SNARE-mediated vesicular exocytosis (Ho, Lee, & Martin, 2011; Sudhof, 2012; Winkle et al., 2014), which in turn can regulate synaptic transmission in the adult brain. The cytoplasmic tail of DCC directly binds to TRIM9, which is a binding partner of the t-SNARE, SNAP25 (Li, Chin, Weigel, & Li, 2001; Plooster et al., 2017; Winkle et al., 2014). Consequently, deletion of DCC may disrupt t-SNARE function to reduce vesicular loading and exocytosis. Although we failed to observe a significant alteration in the mean amplitude of sEPSC events, spontaneous and evoked vesicular pools are regulated differently (Fredj & Burrone, 2009; Melom, Akbergenova, Gavornik, & Littleton, 2013). Fast vesicular exocytosis during evoked stimulation requires simultaneous activation of multiple SNARE complexes, whereas spontaneous exocytosis is thought to depend on a reduced stochiometric requirement of SNARE complex activation (Mohrmann, de Wit, Verhage, Neher, & Sorensen, 2010; Peled, Newman, & Isacoff, 2014). The lack of DCC may impair TRIM9-mediated vesicular release by compromising t-SNARE activation, which may reduce the number of vesicles docked and released from the presynaptic terminal (Plooster et al., 2017; Winkle et al., 2014). Consistent with this hypothesis, we detected a substantial reduction in the amplitude of evoked EPSCs in mice lacking presynaptic DCC at the Schaffer collateral synapse, but no changes in spontaneous fusion, which is thought to engage a distinct vesicular pool and not rely on a synaptotagmin-1-mediated, Ca^2+^-dependent synchronous release mechanism (Acuna et al., 2014; Geppert et al., 1994). While still unclear, DCC association with TRIM9 may mediate vesicular fusion events to gate presynaptic exocytosis of glutamate, and impact synapse function and plasticity (Emptage et al., 2003). Indeed, reduced evoked transmitter release may decrease the ability of Schaffer collateral synapses to initiate post-synaptic depolarization and activate NMDARs, resulting in the attenuation of LTP.

Postsynaptic modification of AMPARs plays a critical role in the expression of HFS-induced LTP, yet activity can also induce changes in both the volume and composition of the presynaptic compartment at Schaffer collateral synapses (Meyer, Bonhoeffer, & Scheuss, 2014). Indeed, presynaptic boutons exhibit enlargement comparable to that observed in postsynaptic spines following plasticity induction, and this enlargement coincides with an increase in the number of synaptic vesicles. Enlargement of presynaptic boutons can occur following glutamate photo-uncaging, suggesting a possible retrograde postsynaptic signal to the presynaptic terminal (Meyer et al., 2014). We have previously demonstrated release of netrin-1 in response to NMDAR activation to facilitate HFS-induced LTP (Glasgow et al., 2018). Together, these findings suggest that HFS may trigger netrin-1 secretion to promote co-ordinated DCC-dependent pre- and post-synaptic changes that underlie the expression of LTP.

### DCC and spatial memory consolidation

Repetitive training can promote dendritic spine formation and clustering *in vivo* (Fu, Yu, Lu, & Zuo, 2012; Peters, Chen, & Komiyama, 2014). Consistent with a role governing dendritic spine dynamics mediated by learning, DCC signaling can potently regulate F-actin cytoskeletal remodeling during development, and may serve a similar function at dendritic spines in the mature nervous system (Koleske, 2013; Lai Wing Sun et al., 2011). Although the mechanisms underlying these changes in spine formation remain unclear, DCC can activate PLC-mediated L-type Ca^2+^ and canonical transient receptor (TRPC) channels to increase intracellular calcium concentration, which critically regulates spine formation (Higley & Sabatini, 2012; Hong, Nishiyama, Henley, Tessier-Lavigne, & Poo, 2000; Wang & Poo, 2005; Xie et al., 2006). Activation of both L-type Ca^2+^ channels and TRPCs have been implicated in synaptic plasticity underlying spatial memory, and genetic deletion of these channels can alter performance on hippocampal-dependent memory tasks (Broker-Lai et al., 2017; Fortin et al., 2012; Marschallinger et al., 2015; McKinney, Sze, Lee, & Murphy, 2009). Deletion of postsynaptic DCC may limit or occlude netrin-1 mediated activation of L-type Ca^2+^ channels and TRPCs to prevent Ca^2+^-mediated *de novo* spine formation or modification of existing dendritic spines. In contrast, deletion of DCC from CA3 neurons may limit Schaffer collateral synaptic plasticity underlying memory formation, resulting in contextual spatial memory deficits. Consequently, conditional deletion of DCC from either CA1 or CA3 pyramidal neurons, which make essential contributions to spatial cognition, may impair structural plasticity and dendritic spine maturation associated with learning and memory formation required for spatial memory tasks, such as the NOPR and Barnes maze.

## Materials and Methods

### Animals

All procedures were performed in accordance with the Canadian Council on Animal Care guidelines for the use of animals in research and approved by the Montreal Neurological Institute Animal Care Committee. R4ag11 (CaMKIIα)-Cre (Dragatsis & Zeitlin, 2000) and Grik4-Cre (Nakazawa et al., 2002) mice were obtained from Jackson Laboratory (Bar Harbor, ME, USA) and maintained on a C57BL/6 genetic background. Both lines were crossed with mice homozygous for floxed *dcc* allele, DCC^*fl/fl*^) (Horn et al., 2013; Krimpenfort et al., 2012). *Cre* recombinase is detectable at P17 and P14 in R4ag11-Cre and Grik4-Cre, respectively. Significant reduction of DCC protein levels in R4ag11 CaMKIIα-Cre-DCC^*f/f*^ (CA1 DCC cKO) and Grik4-Cre-DCC^*f/f*^ (CA3 DCC cKO) mice was observed by 6 months of age, therefore all experiments were performed with mice at least 6 months old. Both males and females were used in behavioural, electrophysiological, and immunohistochemical experiments. We observed no statistically significant differences between sexes, and therefore all data were pooled for analysis. Control experiments were performed using littermates that were negative for *Cre* and homozygous for floxed alleles of *dcc*.

### Western Blots

Animals were initially sedated with isoflurane gas and sacrificed via CO_2_ asphyxiation. Microdissected CA1 and CA3 subregional hippocampal homogenates were collected for Western blot analysis. Tissue samples were homogenized in RIPA buffer supplemented with protease and phosphatase inhibitors (1 mg/mL aprotonin, 1 mg/mL leupeptin, 100 mM PMSF, and 0.5 M EDTA, 1 mM Na_3_VO_4_ and 1 mM NaF). Protein levels were quantified using the bicinchoninic acid protein assay (Pierce BCA kit, Thermo Fisher Scientific) and equal protein concentrations were loaded on acrylamide gels. Proteins were separated using SDS-PAGE electrophoresis on 10% polyacrylamide gels, electroblotted onto nitrocellulose membranes, blocked in 5% non-fat milk in PBS and incubated with primary antibodies overnight at 4°C. The primary antibody used was goat polyclonal anti-DCC A20 (1:1000; Santa Cruz Biotechnology, RRID: AB_2245770). Horseradish peroxidase (HRP) conjugated secondary antibody (1:10 000) was subsequently applied and visualized by reaction with chemiluminesence reagent (Clarity Western Blotting Substrates Kit, BioRad) on radiography film (Carestream Blue X-ray film).

### Barnes Maze

Spatial memory was evaluated in the Barnes maze as described previously (Attar et al., 2013), which consisted of a round table 70 cm in diameter with 12 equally spaced holes along the entire perimeter. One of the holes was designated as the target hole containing an underneath ramp attached to an escape tunnel. Training and test sessions lasted a total of 4 days (Attar et al., 2013; Barnes, 1979). All objects and spatial cues in the room, along with the position of the experimenter, were unchanged during the training and testing sessions. The maze and escape tunnel were cleaned with 70% ethanol between trials to remove any olfactory cues. Mice were trained for 5-6 trials across 2-3 days, with each training trial 4 hrs apart per day. Mice were initially placed in the middle of the table within an opaque starting chamber for 20 secs to randomize the direction they are facing. Once the chamber was lifted, a loud buzzer sound provided motivation to search for the escape tunnel. Each mouse was given 2 mins to search for the target hole independently and allowed to stay in the box for 1 min. The buzzer sound was immediately stopped once the mouse entered the box. For all training sessions, number of nose pokes in the target holes, total number of holes searched, and escape latencies to the escape tunnel were counted by an experimenter blind to the genotype. Twenty-four hrs following the last training session, spatial memory was assessed using a probe trial in which the escape tunnel was removed. For quantitative analysis, the maze was divided into 4 quadrants with 3 holes per quadrants. Quadrants were labelled as: Target, Left, Right, and Opposite, with the target hole being the middle hole in the “Target” quadrant. The amount of time the mice spent in each quadrant was quantified by an experimenter blind to the genotypes.

### Novel Object Placement Recognition

Mice were trained and tested using the novel object placement recognition (NOPR) test as a non-invasive measure of hippocampal-dependent spatial memory, which lasted a total of 3 days (Boyce et al., 2016; Ennaceur, Michalikova, Bradford, & Ahmed, 2005; Wong et al., 2019). On Day 1, mice were habituated to the square testing chamber (50 cm X 36 cm, 26 cm-high wall) for 5 mins without any added objects. On Day 2, during the “Sample Phase”, mice were exposed to two identical objects for 5 mins in two separate training sessions that took place 4 hrs apart. Twenty-four hrs following the last training session called the “Choice Phase”, one of the objects was moved to a novel location in the square chamber and the mice were provided 5 mins to explore both objects. An overhead camera recorded the mice in the square chamber throughout training and testing, and exploration time was measured by an experimenter blind to the genotypes. Objects and the test chamber were cleaned with 70% ethanol between trials to remove any olfactory cues.

Exploration times were calculated as the total time in the probe trial the animal investigated both objects (within one body length from an object with head pointed toward the object). Investigation ratios were calculated as the time spent exploring the novel placed object divided by the total time spent exploring both objects during the probe trial.

### Novel Object Recognition Test

To assess for recognition memory, we employed the novel object recognition (NOR) test (Bevins & Besheer, 2006). The test uses the same protocol as the NOPR test, but rather than moving one of the identical objects to a new location during the “Choice Phase”, one of the objects was replaced with a novel, non-identical object. The test was scored using the same parameters as the NOPR test: total exploration time and investigation ratio.

### Golgi Staining and Spine Morphology

Mouse brains were processed (FD Rapid GolgiStain Kit; FD Neurotechnologies) and cut into 100 μm sections with a cryostat. The length, width, and density of dendritic spines from apical dendrites of CA1 pyramidal neurons were traced and reconstructed, and analyzed using Fiji image software (Schindelin et al., 2012). The type of spine (thin and mushroom) was also recorded.

### *Brain slice* in vitro *electrophysiology*

Acute transverse hippocampal brain slices were obtained from R4ag11-Cre/DCC^*fl/fl*^ (8-10 months old) and Grik4-Cre/DCC^*fl/fl*^ (8-10 months old) mice and age-matched control littermates (Cre-negative DCC^*fl/fl*^). Mice were deeply anaesthetized by intraperitoneal injection of a mixture of 2,2,2 – tribromoethyl alcohol and tert-amyl alcohol diluted at 2.5% in PBS, and transcardially perfused with ice-cold choline chloride-based solution containing (in mM): 110 choline-Cl, 1.25 NaH_2_PO_4_, 25 NaHCO_3_, 7 MgCl_2_, 0.5 CaCl_2_, 2.5 KCl, 7 glucose, 3 pyruvic acid, and 1.3 ascorbic acid, bubbled with carbogen (O_2_ 95%, CO_2_ 5%). The brain was quickly removed and thick horizontal brain slices (250 μM) containing the hippocampus were cut using a vibrating microtome (VT1000s, Leica). Individual brain slices were allowed to recover for 1h in artificial cerebrospinal fluid (ACSF) containing, in mM: 124 NaCl, 5 KCl, 1.25 NaH2PO4, 2 MgSO4, 26 NaHCO3, 2 CaCl, and 10 Glucose saturated with 95% O2 and 5% CO2 (pH ~7.3, 300 mOsm) at room temperature (22°-24° C) prior to recordings.

Individual slices were positioned in a custom-built recording chamber, and continuously perfused with warmed (30 ± 2° C) ACSF (TC324B, Warner Instruments). Current and voltage recordings of CA1 hippocampal pyramidal neurons were performed on an upright microscope (Nikon Eclipse or Scientifica SliceScope 2000) equipped with a micromanipulator (Sutter MP-225 or Scientifica Patchstar), a 40x or 60x water immersion objective (0.8 or 1.0 N.A., respectively), differential interference contrast optics, and coupled to a near-infrared charge-coupled device camera (NC70, MTI or SciCam, Scientifica). Borosilicate glass pipettes (Sutter Instruments) (tip resistance: 4–8MΩ) were prepared using a horizontal puller (P-97, Sutter Instruments), and were filled with an intracellular solution containing (in mM): 120 potassium gluconate, 20 KCl, 10 N-2-hydroxyethylpiperazine N’-2-ethanesulfonic acid (HEPES), 7 phosphocreatine diTris, 2 MgCl_2_, 0.2 ethylene glyco-bis (β-aminoethyl ether)N,N,N’,N’-tetraacetic acid (EGTA), 4 Na^2+^-ATP, 0.3 Na^+^-GTP (pH adjusted to 7.20–7.26 using KOH, 275–85 mOsm). Whole cell somatic recordings from visually-identified CA1 pyramidal neurons were performed in the presence of picrotoxin (100 μM) to block GABAA-mediated inhibitory synaptic transmission. Membrane potential and whole-cell current recordings were obtained using an Axopatch 200B or 700B amplifier (Molecular Devices). Current-clamp recordings were sampled at 20 kHz and filtered at 10 kHz, whereas voltage-clamp recordings were sampled at 10 kHz and filtered at 2 kHz. All data were acquired through an analog-to-digital converter (Digidata 1322A or 1550A, Molecular Devices) for storage on computer hard disk with pClamp software (v9.0 or v10.4, Molecular Devices).

Intrinsic electrophysiological characteristics of CA1 pyramidal neurons were assessed using a series of hyperpolarizing and depolarizing intracellular current pulses (−200 pA to +100 pA). Input resistance was calculated by measuring the peak voltage response to a – 100-pA current step (1000 ms). Rest membrane potential was measured 1 min following break-in to whole-cell configuration.

AMPAR-mediated synaptic currents were evoked using a bipolar platinum/iridium electrode (FHC, CE2C275) placed in the Schaffer collaterals, and placed ~200 μm from the recorded cell. During baseline periods prior to HFS, a pair of 0.1 ms bipolar current pulses (50 ms ISI) was delivered via a stimulus isolation unit (Isoflex, AMPI), and stimulus intensity was adjusted to evoke a response 65-75% of the maximal response at 200 μA. Input-output tests were conducted using increasing stimulus intensity from 0-200 μA in 25 μA increments. Changes in paired-pulse ratio were expressed as the peak amplitude of the second pulse as a function of the amplitude of the first pulse over a range of interstimulus intervals. Access and input resistances were continually monitored throughout the recording through a 50 ms, 5 mV voltage step 150 ms prior to synaptic stimulation, and data were discarded if series resistance changed >20%.

Voltage-clamp recordings at −70 mV were used to measure synaptic transmission at the Schaffer collateral synapse. To assess HFS LTP in both DCC mutants and control littermates, the recording mode of the whole-cell recording was briefly changed to current-clamp, and LTP was induced using a single episode of a 1-s 100 Hz stimulation, which has been previously demonstrated to elicit a long-lasting enhancement of evoked EPSCs in the adult hippocampus. Following stimulation, the cell was returned to voltage-clamp recording mode and held at −70 mV to record evoked responses.

Spontaneous excitatory postsynaptic currents (sEPSCs) were recorded in voltage-clamp mode at a holding potential of −70 mV in the presence of PTX (100 μM) to block GABA_A_-mediated synaptic currents. sEPSCs were analyzed using MiniAnalysis (Synaptosoft), and events were detected using a threshold of 7 pA (>3 pA root mean square of baseline noise levels). Cumulative distribution plots were generated using an equal number of randomly-selected events per condition (100 per condition).

### Statistical Analyses

Statistical analyses on parametric data were assessed using two-way repeated measures analysis of variance (ANOVA) followed by Bonferroni’s post-hoc test, one-way ANOVA followed by Tukey’s pairwise comparison test, and independent *t*-tests where appropriate. Analyses on nonparametric data were assessed using two-tailed Mann-Whitney test. Normality, homoscedasticity, and outlier tests were performed on all datasets. Data were analyzed using Clampfit 10.3 (Axon Instruments), MATLAB (Mathworks), Fiji (Schindelin et al., 2012), Photoshop (Adobe), MiniAnalysis (Synaptosoft), Prism 7 (Graphpad), and Sigmaplot 11 (Systat). Plotted data were then formatted in Adobe Illustrator CS6 (Adobe Systems).

## Author contributions

Conceptualization: SDG, EWW, TEK; Methodology: SDG, EWW, GTS; Investigation: SDG, EWW, GTS; Formal analysis: SDG, EWW, GTS; Writing: SDG, EWW, PS, ESR, TEK; Funding: PS, ESR, TEK; Resources, SDG; Supervision: PS, ESR, TEK

## Acknowledgments

The authors would like to thank members of the Kennedy and Ruthazer labs for comments on earlier drafts of the manuscript. We also thank Nathalie Marcal and Hanna Davies for technical assistance. SDG was supported by postdoctoral fellowships from Fonds de la Recherche Québec – Santé (FRQS) and Canadian Institute for Health Research (CIHR). EWW was supported by a Healthy Brains for Healthy Lives (HBHL) scholarship. The project was supported by grants from CIHR (PS, ESR, TEK). ESR holds a FRQS Research Chair.

